# Reporter gene assays and chromatin-level assays define substantially non-overlapping sets of enhancer sequences

**DOI:** 10.1101/2022.04.21.489091

**Authors:** Daniel Lindhorst, Marc S. Halfon

**Affiliations:** Departments of Biochemistry, University at Buffalo-State University of New York, Buffalo, NY 14203; Biomedical Informatics, University at Buffalo-State University of New York, Buffalo, NY 14203; Biological Sciences, University at Buffalo-State University of New York, Buffalo, NY 14203; NY State Center of Excellence in Bioinformatics & Life Sciences, Buffalo, NY 14203; Department of Molecular and Cellular Biology and Program in Cancer Genetics, Roswell Park Comprehensive Cancer Center, Buffalo, NY 14263

**Author notes:** Correspondence, 955 Main St. #5128, Buffalo, NY 14203.

## Abstract

Transcriptional enhancers are essential for gene regulation, but how these regulatory elements are best defined remains a significant unresolved question. Traditional definitions rely on activity-based criteria such as reporter gene assays, while more recently, biochemical assays based on chromatin-level phenomena such as chromatin accessibility, histone modifications, and localized RNA transcription have gained prominence. We examine here whether these two types of definitions effectively identify the same sets of sequences and find that, concerningly, the overlap between the two groups is strikingly limited. Our results raise important questions as to the appropriateness of both old and new enhancer definitions.

## BACKGROUND

Transcriptional enhancers play an essential role in gene regulation and are primary mediators of development, homeostasis, disease, and evolution [1-3]. Among many unanswered questions about enhancer biology, one stands out as fundamental: how should an enhancer be defined? In the genomic era, the functional definition that for a quarter-century described enhancers as sequences with the ability to drive expression of a reporter gene from a minimal promoter [4, 5] has entered into an uneasy co-existence with transcription and chromatin-based definitions such as ability to bind specific sets of transcription factors or coactivators, presence of certain histone modifications, location in nucleosome-depleted regions, or transcription of enhancer RNA (eRNA) [e.g. 6, 7-15]. It has become increasingly clear that these chromatin-level enhancer definitions identify sets of sequences with strikingly low levels of overlap, with concerning implications for regulatory genome annotation, ongoing studies of enhancer biology, and interpreting results from genome-wide association studies, expression quantitative trait locus studies, and the like [16-18]. However, it is unknown which of these various assays are in the best agreement with reporter gene data, which remains the “gold standard” for enhancer activity, and to what extent. Here, we perform a comprehensive comparison to investigate whether one or a collection of chromatin-based assays are able to identify the majority of enhancers from an extensive reporter-gene defined set. We show that not only do the chromatin-level assays show poor agreement among themselves, but also that they fail to discover a significant fraction of reporter-gene defined enhancers, often performing no better than random expectation. Our results raise questions as to whether any common current assays sufficiently interrogate the enhancer landscape, and about the accuracy of current regulatory genome annotations.

## RESULTS AND DISCUSSION

In order to test for congruence between enhancers defined by reporter gene assays, and those defined by chromatin-based assays, we compared *Drosophila* enhancers in the REDfly database [19] with those in the EnhancerAtlas2.0 database, which uses a supervised learning approach to combine data from ChIP-seq, ATAC-seq, FAIRE-seq, and other chromatin-level assays [20]. This is uniquely possible for *Drosophila*, as close to 16,000 of REDfly’s >33,000 enhancers are drawn from reporter gene assays based on small-scale “enhancer-bashing” studies and therefore not confounded with the 294,158 predicted *Drosophila* enhancers in EnhancerAtlas.

For an initial comparison, we took all REDfly enhancers 2 kb or shorter (11,549 total) and determined how many of these sequences overlapped an EnhancerAtlas enhancer (see Methods). Of the 21 tissue-specific *Drosophila* EnhancerAtlas datasets, 17 had statistically significant overlap (adjusted *P* < 0.01, two-tailed *z* test), three had no significant overlap, and, surprisingly, one had significantly less overlap than expected by chance (Fig. 1A “STARR-seq included”, Table S1a). Examination of these data revealed that more than three-quarters of the REDfly enhancers (8779/11549, 76%) were identified using a single method, STARR-seq [21], and that the STARR-seq enhancers made up 80% of the REDfly-EnhancerAtlas overlapping enhancers (6027/7527). When we eliminated the STARR-seq enhancers from the REDfly test set and repeated the analysis, the results were strikingly different: only five of the EnhancerAtlas datasets (24%) now showed significant overlap (with one additional dataset just below our significance threshold), whereas ten datasets (48%) had significantly less overlap than expected by chance (Fig. 1A “STARR-seq not included”, Table S1b).

**Figure 1:**
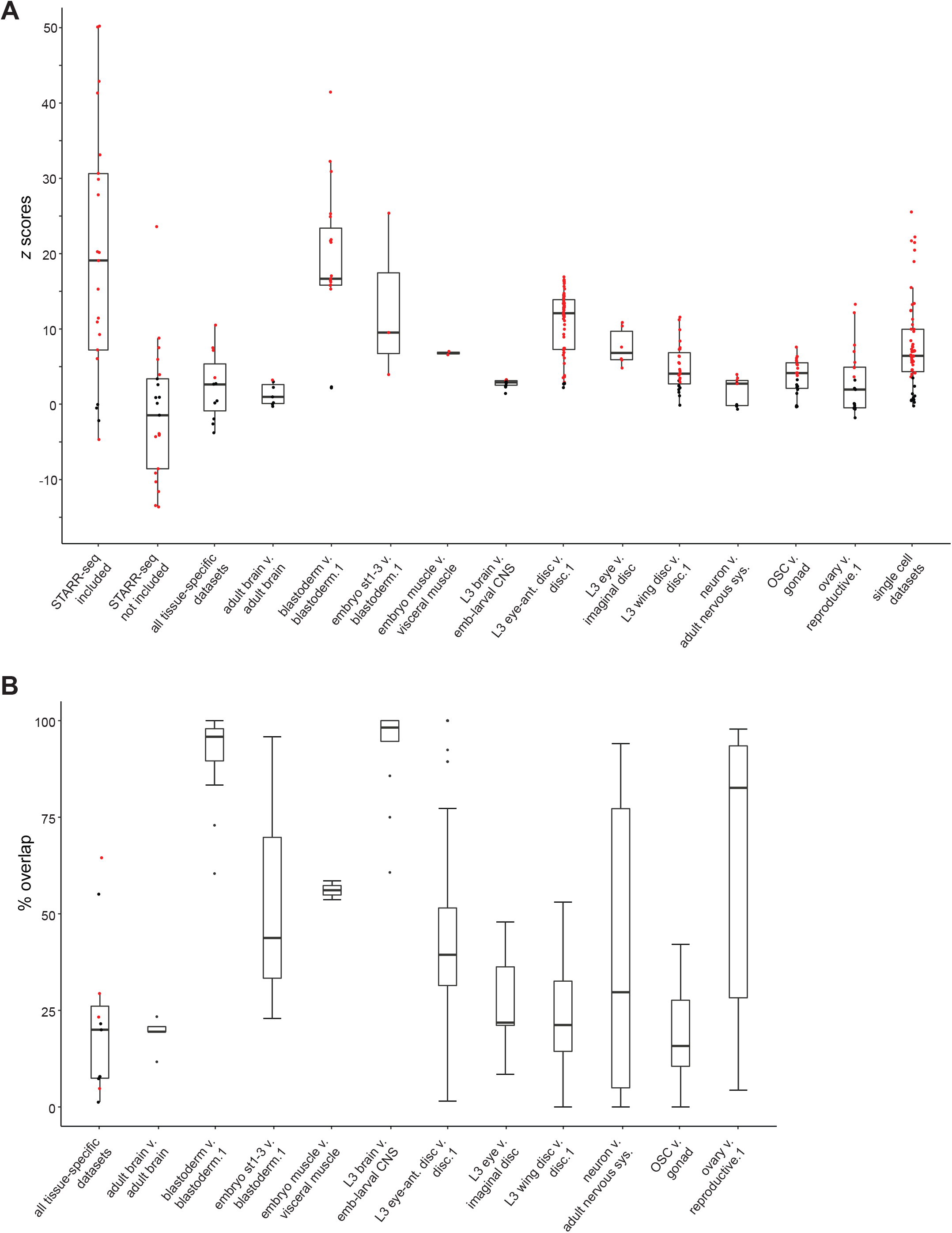
Comparisons between REDfly and EnhancerAtlas enhancer sets. Box plots show medians and the first and third quartiles. Data points shown in red are significant at a Bonferonni-adjusted *P*-value < 0.01. Dataset names correspond to the names in Table S1. For the “tissue-specific comparisons” (Table S1c), when multiple REDfly sets corresponded to the same EnhancerAtlas set, the one with the highest degree of overlap was selected for analysis. For the subset comparisons (Table S1d,e), the REDfly set with the largest number of significantly overlapping individual EnhancerAtlas component sets was used. (A) *z*-score distributions for each of the REDfly-EnhancerAtlas comparisons. (B) Percent of REDfly enhancers found that overlap EnhancerAtlas enhancers. For the “all tissue-specific datasets” plot, datasets with significant *z*-scores are indicated in red. For the subset comparisons, values represent the median percent overlap of the constituent data subsets, as shown in Table S1e.

STARR-seq-defined enhancers thus have a profound effect on how well sequences from the two databases compare. Only four EnhancerAtlas datasets make use of STARR-seq data, indicating that a simple confounding of the source data cannot explain the results. In the absence of STARR-seq enhancers, almost half of the REDfly datasets had less-than-expected overlap with EnhancerAtlas, suggesting that the majority of REDfly enhancers lack activity in most of the tissues covered by EnhancerAtlas. Conversely, these results suggest that STARR-seq, performed in a cultured cell line, may be identifying many sequences indiscriminately with respect to their tissue specific activity.

To examine more directly how REDfly enhancers active in specific tissues compare with the EnhancerAtlas datasets, we constructed tissue-specific REDfly enhancer sets using only in-vivo reporter gene tested enhancers < 1000 bp in length (Table S2). Of the 21 EnhancerAtlas sets, 11 had one or more corresponding REDfly sets. We then determined how many enhancers from each paired set overlapped. Our expectation was that there should be significant overlap, with the EnhancerAtlas sets (based on undirected genome-wide assays) encompassing the great majority of REDfly enhancers (drawn from individual *ad hoc* experiments). Surprisingly, only four (36%) of the eleven EnhancerAtlas sets showed significant overlap with their corresponding REDfly set (*P* < 0.01, Fig. 1A “all tissue-specific datasets”, Table S1c). Moreover, even for the sets with significant overlap, the number of in-common enhancers was strikingly limited (median 26%, range 5%-65%; Fig. 1B, Table S1c).

Since the EnhancerAtlas definitions integrate the data from multiple assays, we reasoned that the integration algorithm might be filtering out some of the true enhancers. To test this, we took the underlying data sets used by EnhancerAtlas and compared them individually to the matched REDfly enhancer sets. Indeed, we saw a higher number of significantly overlapping enhancer sets (66%, Fig. 1A, Table 1d,e), but again, the number of in-common enhancers within each matched set was low (median 39%, Fig. 1B, Table 1d,e).

The low degree of in-common enhancers could represent a small number of REDfly enhancers that consistently match EnhancerAtlas enhancers, or different subsets of REDfly enhancers for each EnhancerAtlas subset. To distinguish between these possibilities, we looked at the correlation between the sets of REDfly enhancers present in individual EnhancerAtlas subsets. The sets of enhancers found in different experiments were not overall well-correlated, suggesting that distinct REDfly enhancer sets are being identified (Fig. 2, Fig. S1). However, a clear correlation structure was evident in which assays from a particular laboratory and method tended to cluster together. For example, the REDfly enhancers from series GSE102839 are highly correlated (Fig. 2, white box; mean *r* = 0.74 ± 0.11), and those from series GSE101827 are highly correlated (Fig. 2, yellow box; mean *r* = 0.67 ± 0.18), but these two sets of ATAC-seq experiments find a disjoint group of REDfly enhancers (mean *r* = 0.12 ± 0.08). Although assay-specific correlation is occasionally observed (Fig. 2, yellow asterisks; *r* = 0.77), in other cases similar assays are only weakly correlated (e.g., Fig. 2, white asterisks; *r* = 0.45). EnhancerAtlas data based on ATAC-seq assays had a moderate bias toward significant REDfly overlap (18/26 significant; data not shown), while those based on ChIP directed against RNA pol2 tended toward poor overlap (12/31 significant). However, the batch effects dominate the correlation structure, making it difficult to determine the most effective enhancer identification methods. These observations suggest two important conclusions: (1) identification of putative enhancers is highly dependent on not just the type of assay performed, but on the precise conditions under which it is performed; and (2) enhancer identification is reasonably robust given a particular set of assays and replicates. The fact that the sets of REDfly enhancers are stable within groups of replicate assays suggests that these enhancers are being specifically (i.e., non-randomly) found, despite the fact that, as shown above, the number of REDfly enhancers identified through chromatin-level assays is frequently indistinguishable from random expectation. Thus, these assays do appear to be able to identify reporter-gene-defined enhancers, but with low efficiency and potentially high false-positive rates.

**Figure 2:**
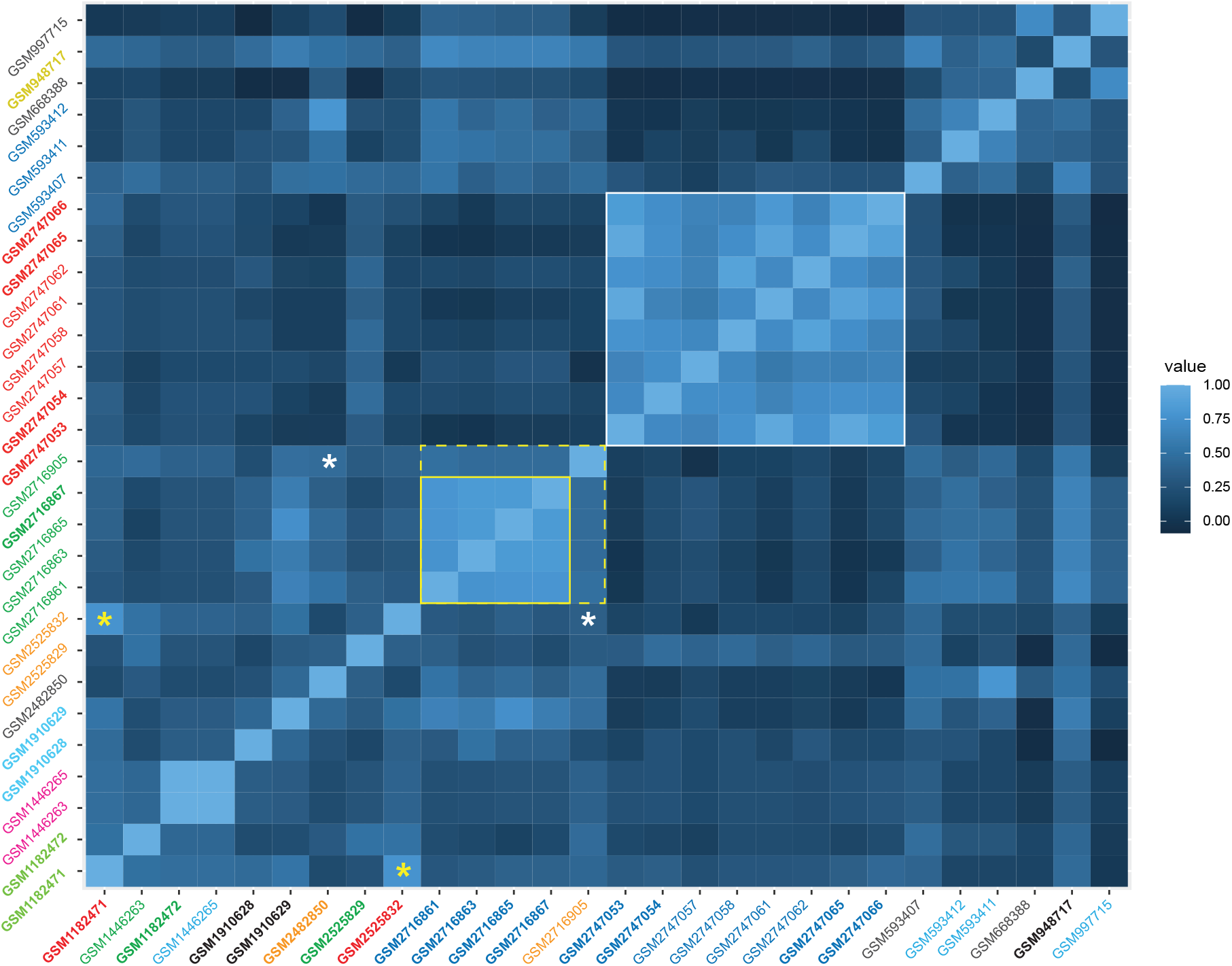
Correlations between REDfly enhancers overlapping enhancers from each EnhancerAtlas subset used for the “L3_wing_disc” EnhancerAtlas set. The correlation structure demonstrates that experimental batch effects predominate over assay-type effects. High correlations are seen between sets from the same experiment group (see white box, yellow box), even when assay types differ (dotted yellow box). Although assay-specific correlations are sometimes present (yellow asterisks), identical assays can also be poorly correlated when from different experiment groups (white asterisks). Label colors along the y-axis represent common experiment series as provided in GEO (see Table S1d). Sets labeled in black are unrelated. Label colors on the x-axis indicate assay types as follows: red, H3K4me1 ChIP-seq; green, H3K27ac ChIP-seq; orange, Grh ChIP-seq; cyan, Pol2 ChIP-seq; dark blue, ATAC-seq; black, ChIP-seq against various transcription factors.

Recently, Gao et al. released “scEnhancer”, an EnhancerAtlas-like database based on single-cell ATAC-seq data [22]. Unlike the results using EnhancerAtlas, the majority of scEnhancer datasets had a significant degree of overlap with their REDfly set (45/56, 80%; Fig. 1A, Table S1f), although similar to what we observed with EnhancerAtlas, the number of overlapping enhancers was low (median 27%, range 10%-65%; Table S1f). scATAC-seq may thus represent a more promising method for enhancer detection, although a proper assessment is difficult as all of the *Drosophila* data currently in scEnhancer are from a single source.

## CONCLUSIONS

Reporter gene assays have long been considered the gold standard for defining enhancers. On these grounds, our results would seem to suggest that not only do chromatin-level assays frequently fail to identify common sets of enhancer sequences [16], but neither are they particularly effective at covering the majority of the enhancer landscape. However, we would caution against automatically accepting this conclusion, as there are well-known deficiencies that could lead to a substantial number of both false-positive and false-negative results from reporter gene assays. These include enhancer-promoter incompatibility, effects due to episomal expression or chromosomal integration, cell-type specificity, and ectopic expression due to missing repressor binding sites, as well as recent findings that enhancers can double as silencers or functionally overlap other regulatory features [see discussions in 23, 24-27]. Without true “known” enhancer data, therefore, it is impossible to say whether our results reflect significantly poor sensitivity in chromatin-level assays, or a much larger than heretofore recognized false positive rate in reporter gene assays. What is indisputable, however, is a clear need for approaches that can reconcile the poor agreement between and among the various reporter gene and chromatin-level enhancer identification methods. In this regard, the increasing tractability of genome-engineering approaches, e.g., via CRISPR/Cas9 sequence deletion and replacement, holds out an encouraging potential to interrogate the enhancer capability of sequences within their native genomic contexts.

## METHODS

### Data sources

For comparisons using all REDfly data, sequences were obtained from REDfly v7.1.1 (Aug. 14 2020) by downloading all “CRM” entries in bed format, eliminating sequences > 2000 bp, and then removing overlapping sequences using the script “SelectSmallestFeature.py” (Kazemian and Halfon 2019).

For comparisons using specific REDfly subsets, “CRM” sequences were downloaded from REDfly v5.6.1 (Dec. 3 2019) in bed format with “cell-culture only” sequences excluded and a sequence length cutoff of either 1000 or 600 bp. Following removal of overlapping sequences using “SelectSmallestFeature.py” (Kazemian and Halfon 2019), the expression pattern annotations associated with each remaining sequence were used to place the sequences into one or more of 30 different tissue-specific groupings. Details of the composition of each set can be found in Supplementary Table S2.

For compatibility with EnhancerAtlas, genome coordinates were converted from dm6 to dm3 using LiftOver [28] with minMatch = 0.25.

EnhancerAtlas sequences were downloaded from EnhancerAtlas 2.0 (http://enhanceratlas.org, Nov. 19 2019). Identities of the component datasets for each of the 21 tissue-specific EnhancerAtlas sets were obtained from the metadata files available in the “data source” section. The provided GEO accession codes were then used to obtain the sequence-level data from NCBI. Data processing was performed as described in the EnhancerAtlas paper [20] to ensure consistent results. Wig files were converted to bigWig format through the script wig2BigWig, downloaded from the UCSC genome browser at genome.ucsc.edu/goldenpath/help/bigwig.html. Bigwig files were converted to bedgraph format through the script bigWigtoBedGraph. Bedgraph files were converted to bed format through peak calling using MACS2 [29] with a cutoff enrichment of 2. Data sets where we were unable to replicate the exact EnhancerAtlas processing pipeline (primarily, raw sequencing data) were omitted from further analysis.

scEnhancer sequences were downloaded from scEnhancer (http://enhanceratlas.net/scenhancer/, Feb. 28. 2022).

### Comparison of Data Sets

Bed files were compared using BEDtools *intersect* [30] and the *-wa* and *-u* flags. Note that with these parameters, even a single intersecting basepair will cause the two sequences to be scored as an intersection, making our tests highly sensitive to any degree of sequence overlap.

### Significance of comparisons

Significance of dataset overlap was determined by permuting the coordinates of each REDfly dataset 500 times using BEDtools *shuffle* [30] and repeating the tests for intersection. The mean and standard deviation of the permuted results were then used to calculate a *z*-score. A Bonferroni-corrected *P* value equivalent to *P* < 0.01 was determined for each set of comparisons.

### Correlation analysis

For each comparison, each potential REDfly enhancer was scored as 1 (found) or 0 (not found). Correlations between all pairs of result vectors were determined using the R *cor* function and visualized as heat maps using *ggplot*.

## Authors’ contributions

DL performed the analysis. MSH conceived of the study and provided guidance on the analysis. Both authors wrote the manuscript. Both authors read and approved the final manuscript.

## Acknowledgements

We thank Hasiba Asma for assistance with figures and Yungki Park, Michael Buck, and members of the Halfon lab for comments on the manuscript. Nanzeeba Ahmad performed some preliminary analyses for this project. This work was performed in part at the University at Buffalo’s Center for Computational Research (http://hdl.handle.net/10477/79221).

---

**Figure S1.**
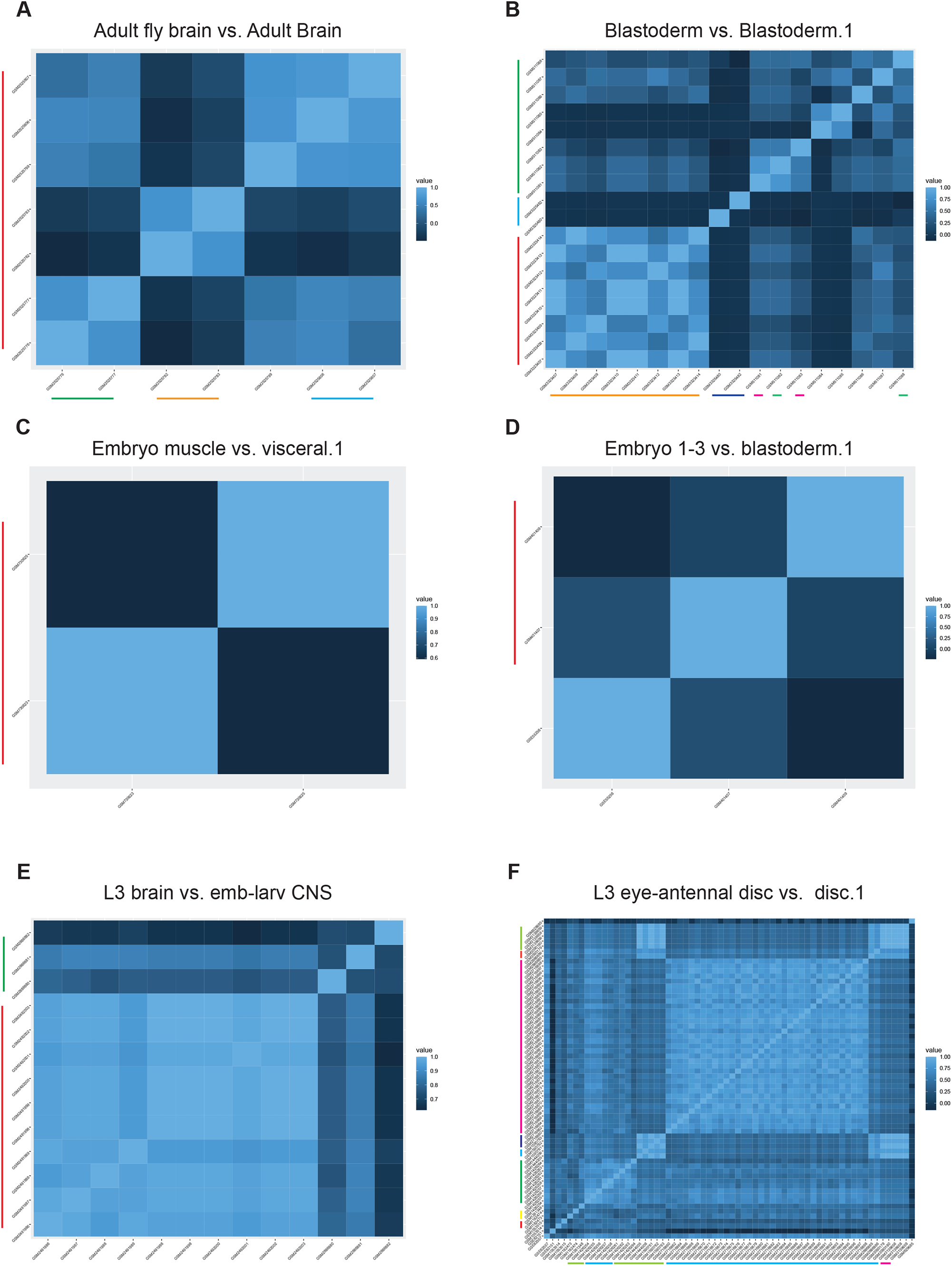
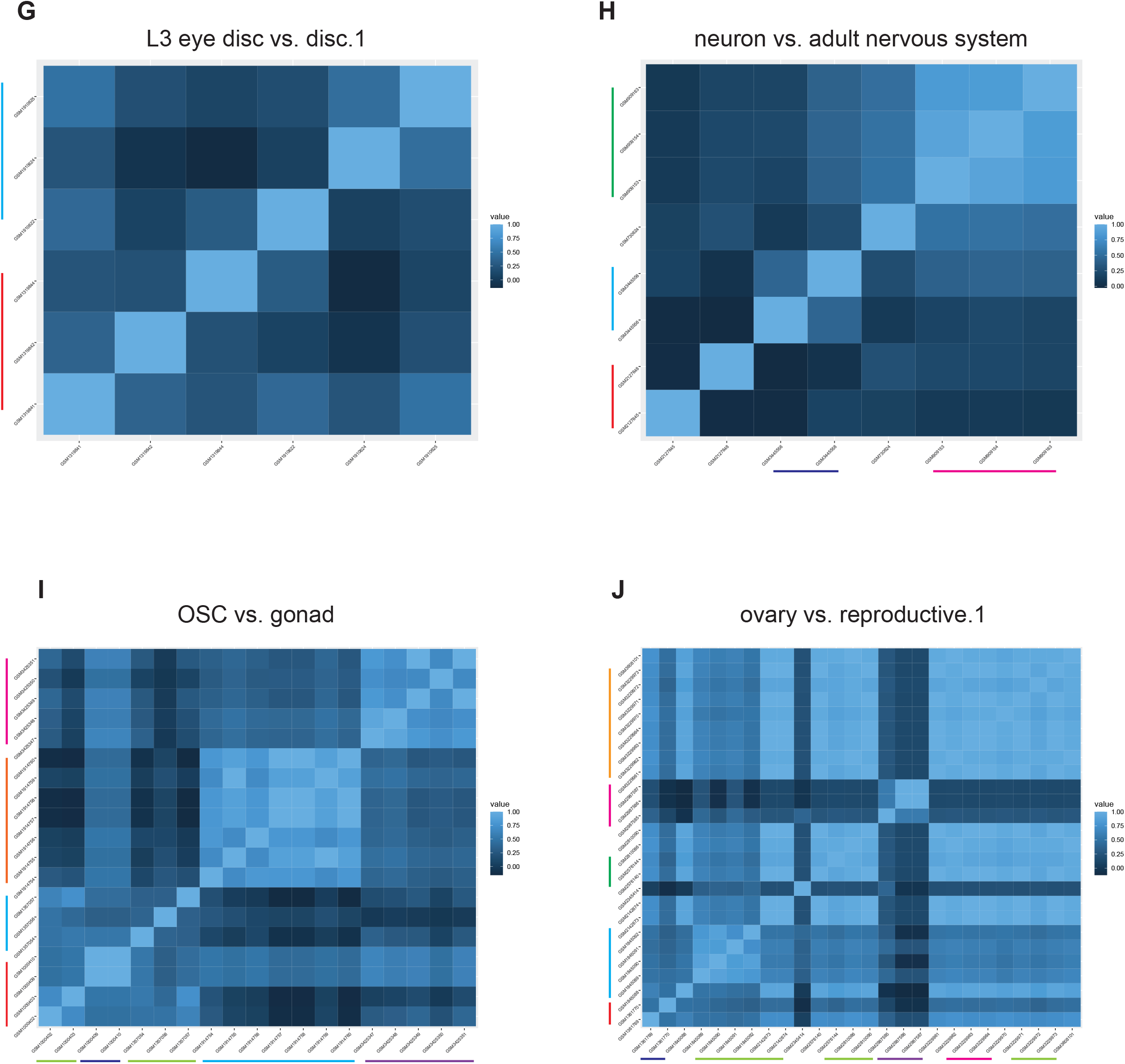
Lindhorst and Halfon 2022.

